# Delay in flowering time in Col-0 under water deficit and in *ddc* triple knockout mutant

**DOI:** 10.1101/2025.03.11.642597

**Authors:** Emil Vatov, Tsanko Gechev

## Abstract

This study addresses the role of cytosine methylation in the fine tuning of flowering time under conditions of water deficit in *Arabidopsis thaliana*. For this purpose, the *drm1 drm2 cmt3* triple methylation mutant in the Col-0 background was used. The plants were grown under long day conditions with water deficit induced by cessation of watering starting at 12 days after seeding. Col-0 showed 1 day delay in flowering as a result of the treatment. *ddc* on the other hand, showed 2 days of delay regardless of the experimental conditions. We found that the two b-box domain proteins BBX16/COL7 and BBX17/*COL8* became overexpressed in the *ddc* background and in Col-0 under water deficit at 24 days after seeding. Additionally, the *NF-YA2* transcription factor became correspondingly down-regulated. Our results suggest a model, where BBX16/COL7 and BBX17/*COL8* interact with CONSTANS to delay the induction of *FT* under long day conditions. *NF-YA2*, which is also recognized as a promoter of *FT* expression, with its down-regulation causes additional delay of *FT* induced flowering. In the Col-0 background, the weak *FRIGIDA* allele fails to induce sufficient expression of *FLC* for additional suppression of *FT*. The plants overcome easily the BBX/NF-YA inhibition resulting in a relatively small delay in flowering. The expression patterns of the three genes suggest involvement of cytosine methylation in their regulation, however no differential methylation could be found in *cis* that can explain these effects. The results therefore, suggest a *trans* acting mechanism. Considering that the activities of *BBX16*/*COL7* and *BBX17*/*COL8* in different physiological conditions are not elucidated, this paper provides a background for future experiments targeting the role of these genes in the fine tuning of flowering time in *Arabidopsis thaliana*.

## Introduction

Stress induced disturbances in flowering time are of increasing importance, not only due to the lack of irrigation water in many parts of the world, but also due to global climate changes leading to sporadic hot and dry spells. Better understanding of the molecular mechanisms underlying flowering time under drought can help breeders and farmers utilize varieties better suited for their conditions. For example, a region experiencing long dry spells may benefit from varieties that induce earlier flowering and avoid the stress altogether. On the other hand, areas with relatively short and sporadic dry spells may benefit from varieties that can inhibit their growth and flowering in order to endure the stress and continue development. In this study, the focus will be placed on the later strategy of stress tolerance and inhibition of flowering time as a response to water deficit.

The initiation of reproduction in plants happens on the basis of large amounts of internal and external cues (Freytes et al., 2021; Hyun et al., 2017; Sharma et al., 2020; Wu et al., 2020). Two types of strategies are recognized in *Arabidopsis* in relation to flowering time under drought stress – avoidance and tolerance. Lovell et al., (2013) provided evidence that natural variation in the expression of *FRI* (*FRIGIDA*) correlates well with physiological parameters related to the two strategies. A low expression allele confers a “drought escape” strategy with fast growth, low water use efficiency and early flowering, while “dehydration avoidance” is related to slow growth, efficient water use and late flowering (Lovell et al., 2013). Utilizing a QTL population between Landsberg *erect* and Antwerp-1 Schmalenbach et al., (2014) confirmed that early flowering as a drought escape strategy is related to strongly impaired plant fitness, while late flowering plants were able to recover their growth in the second half of their vegetative development. Indeed, stress tolerance in the face of late flowering may be advantageous under conditions of mild water deficit (Schmalenbach et al., 2014). Col-0, the wild type used as a control in the present study, is characterized as a “drought escape” phenotype with impairment in the *FRI* allele (Lovell et al., 2013). As such, it provides a good background for studying additional pathways that act to delay flowering time, which may be masked behind a strong *FRI* allele.

*FRI* is known to regulate flowering by promoting the expression of *FLOWERING LOCUS C* (*FLC*), via establishment of active chromatin states (Li et al., 2018). *FLC* can then suppress the expression of *FLOWERING T* (*FT*) and *SUPPRESSOR OF OVEREXPRESSION OF CO 1* (*SOC1*). While Col-0 contains a weak *FRI* allele, there are other pathways that converge on the *FLC* locus. For example, some evidence exists that the *miRNA169* regulated *NF-YA2* can promote the expression of *FLC* (Xu et al., 2014). On the other hand, *NF-YA2* was also shown as a positive regulator of *FT* (Siriwardana et a., 2016; Gyula et al., 2018). It is odd that the same transcription factor that affects positively the main gene responsible for induction of reproductive growth will also affect positively its main suppressor. Recent reviews also appear to have conflicting opinions regarding the role of *NF-YA2* on flowering. Siriwardana, (2024) maintains the positive effects of the *NF*-*Y* complex on flowering, without mentioning the impact of *NF-YA2* on *FLC*, while Chen et al., (2023) depicts a model where *NF-YA2* promotes the expression of *FLC*, which in turn suppresses *FT*, without mentioning its effects on *FT*.

In another aspect of *FT* regulation, the B-box domain transcription factor *CONSTANS* (*CO*) is recognized as the main trigger of *FT* expression under long-day conditions (An et al., 2004). Recently, two members of the B-box family *CONSTANS*-*LIKE 7* (*BBOX16*/*COL7*) and *CONSTANS*-*LIKE 8* (*BBOX17*/*COL8*), were found to interact directly with *CO* and inhibit its function as *FT* promoter under long day conditions (Susila et al., 2023; Xu et al., 2022). The present study provides evidence that *COL7* and *COL8* become overexpressed under water deficit conditions, in a cytosine methylation dependent manner, and may be responsible for the delay of flowering time observed in Col-0.

For the purpose of this study, the *drm1 drm2 cmt3* triple methylation mutant in the Col-0 background was used. The *DOMAINS REARANGED METHYLASE 1*/*2* (*DRM1*/*2*) are recognized as the main methyltransferases participating in the RNA-directed DNA methylation pathway in *Arabidopsis* (Cao and Jacobsen, 2002a; Cao and Jacobsen 2002b). In this context, the two enzymes are primarily responsible for cytosine methylation in euchromatic regions. On the other hand, *CHROMOMETHYLASE 3* (*CMT3*) acts together with proteins from the SUVH family to form a self-reinforcing feedback loop with the histone modification H3K9me (Du et al., 2012). In this action, *CMT3* is mainly responsible for cytosine methylation in CHG and CHH contexts, where H is any nucleotide except G. Additionally, *Arabidopsis* has the *MET1* methyltransferase, which facilitates CG methylation maintenance in interaction with *VIM1* and H3K9me2/3 (Kim et al., 2014). Also, the *CMT2* methyltransferase, which together with *CMT3* facilitate CHG/CHH methylation (Du et al., 2012), and together with *DECREASED DNA METHYLAITON 1* (*DDM1*) facilitate methylation in heterochromatic regions (Zemach et al., 2013). Altogether, the *ddc* tripple knockout mutant is characterized by an almost complete loss of CHG methylation, severe reduction in CHH and mild reductions in CG methylation (Stroud et al., 2013). Additionally, the *ddc* was previously reported to exhibit a slight delay in flowering time under long day conditions (Vatov et al., 2022).

This study was performed under the hypothesis that if certain cytosine methylation patterns, established by the *DRM1*/*2* and *CMT3* methyltransferases and lost during water deficit, play an important role in the regulation of flowering time under water deficit in Col-0, the *ddc* mutant should not exhibit additional delays. Furthermore, differentially regulated genes responsible for the regulation of flowering time under water deficit should already be differentially expressed in the *ddc* mutant under control conditions and should share the same patterns of expression. If differential methylation is the main prerequisite for differential regulation under water deficit, then no differential expression is expected in the *ddc* mutant under stress. Here we show that, indeed the

*COL7* and *COL8* b-box domain proteins meet these criteria, together with the *NF-YA2* transcription factor. Surprisingly, however, we did not find any changes in the cytosine methylation landscape within, or around the gene bodies, which could explain the changes in expression, indicating a *trans* regulatory mechanism. Considering that the activities of *COL7* and *COL8* in different physiological conditions are not elucidated, this paper provides a background for future experiments targeting the role of these genes in the fine tuning of flowering time in *Arabidopsis thaliana*.

## Materials and methods

### Plant growth

The seeds for this experiment were obtained from the Nottingham Arabidopsis Stock Center - NASC. The *ddc* triple knockout mutant (NASC ID: N16384) was grown for one generation for seed production and then utilized for this experiment. Approx. 5-10 seeds were placed in a pot, filled with a moist mixture of 3 parts peat substrate to 1 part perlite. The seeds were stratified for 48 h in complete darkness at 4 °C. after that the plants were grown under long days (16 h day/8 h night) at 22 oC and 70% relative humidity. after germination the pots were thinned until 1 plant per pot was le*FT*. Plants were grown in a randomized complete block design. Each block consisted of 2 trays next to each other, one of them used as a control and the other as a low watering treatment (Figure 1). The control was watered two times a week. For the treatment, watering was stopped for 12 days starting at day 12 after seeding. The total number of plants used was 160. At 24 days after seeding (DAS), before watering, 96 plants were harvested for further analysis. Bolting and rosette diameter of all plants was estimated visually. Bolting was evaluated in DAS when the shoot apical meristem has visibly transitioned from vegetative to reproductive growth. Rosette diameter was evaluated from pictures using ImageJ. after harvesting the plants’ rosette fresh weight was estimated. Leaf number five was used for relative water content. The rest of the plant material was frozen in liquid nitrogen and stored at -80 ° C Relative water content

**Figure 1.**
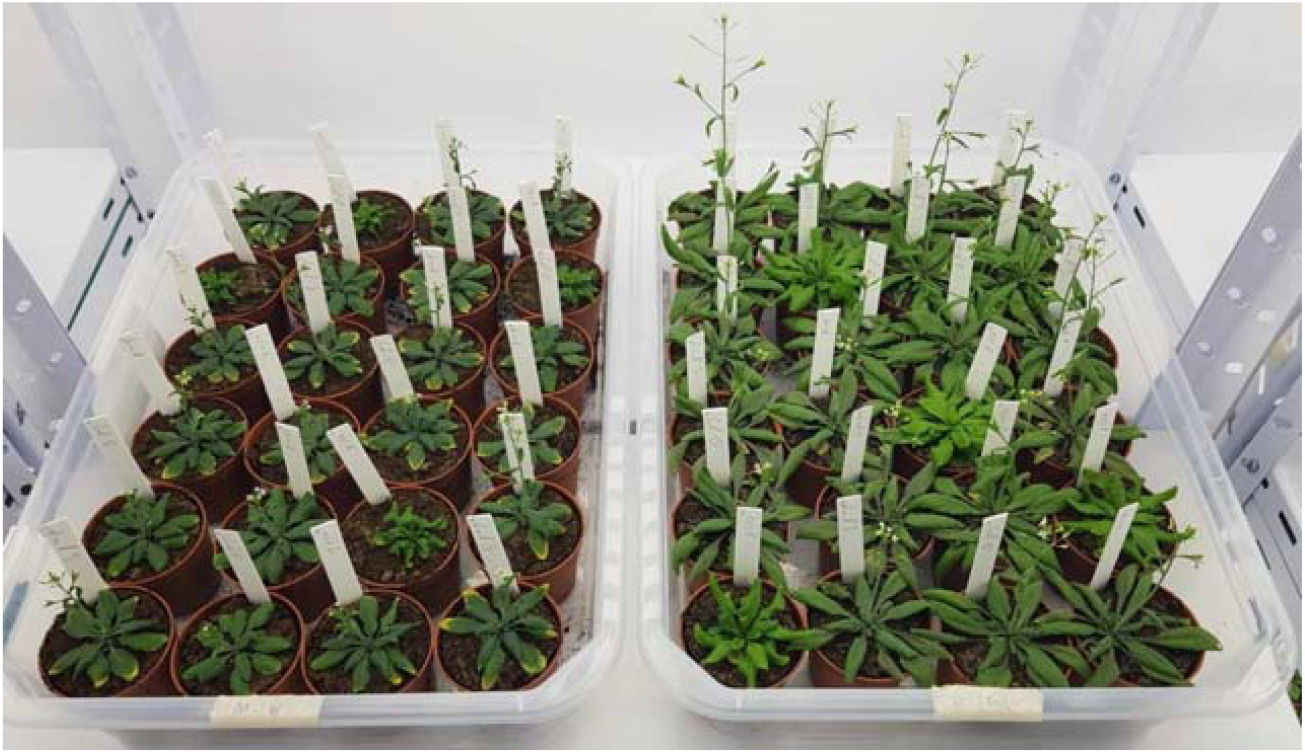
Col-0 and *ddc* during flowering. On the left is the Low Watering treatment, on the right is the Control.

Relative water content from leaf number five was estimated with the following formula: 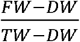

Here, FW is the leaf fresh weight; DW is the leaf dry weight; and TW is the leaf turgid weight after 24 h in distilled water, before the beginning of the dehydration process.

### RNA extraction

For RNA extraction, the 96 plants were pooled in 12 samples and ground in liquid nitrogen. These are the four treatment conditions (Col-0 Control, Col-0 Low Water, *ddc* Control and *ddc* Low Water) with three biological replicates. RNA was extracted with PRImeZOL™ from Canvax according to the manufacturer’s instructions, with slight modifications. 40-50 mg frozen plant material was placed in 1mL of PRImeZOL™ and homogenized for 1 min in a VWR Star-Beater at a frequency of 30/s. The samples were centrifuged at 12000 g for 10 min at 4 °C. The supernatant was transferred to a new tube and incubated for 1 min at room temperature. 200 µl of chloroform were added, the sample was mixed by pipetting up and down carefully and incubated for 3 min at room temperature. after centrifugation at 12000 g for 15 mins at 4 °C, approx. 400 µl of the upper aqueous phase were moved to a new tube. 500 µl of isopropanol were added and the samples were incubated for 10 min at room temperature. Following 10 min of centrifugation at 12000 g and 4° C, the supernatant was carefully removed and two steps of washing with 75% ethanol ensued. Washing was performed with 10 min incubation at 200 rpm and room temperature, followed by centrifugation at 7500g for 5 min at 4 °C and change of the supernatant. after that, the pellet was air dried for approx. 15 mins, or until it loses its whitish color and becomes transparent. The RNA was dissolved in 50 µl of RNAse free water and incubated for 10 min at 60° C.

### RNAseq

The RNA from the 12 samples was sequenced by BGI Genomics on the DNBSEQ platform with PE150 sequencing read length and DNBSEQ Eukaryotic Strand-specific mRNA library (Supplementary Table 1). The raw data was filtered with SOAPnuke (Chen et al., 2018) with the following parameters: *-n 0*.*001 -l 20 -q 0*.*4 --adaMR 0*.*25 --polyX 50 –minReadLen 150*. The read quality was checked using FastQC (Andrews 2017). Genome indexing and alignment were perfirmed with STAR (Dobin et al., 2013). The following parameters were used for indexing: *--sjdbOverhang 299 -- genomeSAindexNbases 12*. The following parameters were used for alignment: *--outSAMtype BAM Unsorted --quantTranscriptomeBan Singleend --outFilterType BySJout --alignSJoverhangMin 8 -- outFilterMultimapNmax 20 --alignSJDBoverhangMin 1 --outFilterMismatchNmax 999 -- outFilterMismatchNoverReadLmax 0*.*04 --alignIntronMin 20 --alignIntronMax 6000 -- alignMatesGapMax 1000000 --quantMode TranscriptomeSAM --outSAMattributes NH HI AS NM MD*. Quantification was then performed using Salmon (Patro et al., 2017). The TAIR10 genome assembly was used for this analysis.

### DNA extraction

DNA was extracted from the same 12 samples used for RNAseq. Approx. 100 mg of plant material were placed in 500 µl CTAB extraction buffer (2% cetyl trimethylammonium bromide, 1% polyvinylpyrrolidone, 100 mM Tris-HCl, 1.4 M NaCl, 20 mM EDTA), vortexed thoroughly and incubated at 60 °C for 30 minutes. after centrifugation for 5 mins at 14000 g, the supernatant was transferred to a new tube and 5 µl of RNAse A solution was added. The samples were incubated for 20 min at 37 °C. after that, 500 µl of phenol:chloroform:isoamyl alcohol (25:24:1) were added and mixed by pipetting. The phases were separated via centrifugation for 1 min at 14000 g. The upper aqueous phase was moved to a new tube and 0.7 volumes of cold isopropanol were added. Precipitation was enhanced at -20°C for 15 min and DNA was pelleted at 14000 g for 10 min. The isopropanol was removed and two subsequent washing steps with 75% ethanol were performed, with 10 min incubation time for each. The DNA pellet was air dried for approx. 15-20 min and dissolved in 50 µl TE buffer (10 mM Tris, pH 8, 1 mM EDTA).

### Whole genome bisulfite sequencing

The DNA from the 12 samples was sequenced by BGI Genomics on the DNBSEQ platform with PE150 sequencing read length and DNBSeq Whole genome bisulfite library (Supplementary Table 2). The raw data was filtered with SOAPnuke (Chen et al., 2018) with the following parameters: *-n 0*.*001 -l 20 -q 0*.*4 --adaMR 0*.*25 --ada_trim –minReadLen 150*. The read quality was checked using FastQC (Andrews 2017). Sequence alignment was performed using Bismark (Krueger and Andrews, 2011) with the following settings: *--multicore 8 --score_min L,0,-0*.*4*. The alignments were then sorted with Samtools (Danecek et al., 2021). The data was deduplicated and cytosine methylation information was extracted with Bismark. Methylation extraction was performed with the following settings: *-p -- ignore 3 --ignore_r2 3 --comprehensive --bedGraph --CX --multicore 10*. The TAIR10 genome assembly was used for this analysis.

### Statistical analyses

All statistical analyses and data visualizations were performed on R 4.3.1. Rosette fresh weight, rosette diameter, relative water content, bolting and global methylation levels in CG, CHG and CHH were evaluated using a mixed linear model. Treatment, genotype and their interaction were used as main effects, while the block structure of the experiment was used as a random effect. This was done with the help of the *lme4* package (Bates et al., 2014). Estimated marginal means and contrasts were calculated with the *emmeans* package (Lenth, 2024). The data was visualized with the help of *ggplot2* (Wickham, 2016), ggpubr (Kassambra, 2023) and *cowplot* (Wilke, 2024).

Statistical analysis of the RNAseq data was performed using the *DEseq2* pipeline (Love et al., 2014). All genes with less than 10 reads in more than 9 samples were removed from the analysis. PCA was performed and visualized with the *plotPCA* function from the *DEseq2* package. Differential expression was calculated, accounting for the block effect of the experimental design. Volcano plots were made using *ggplot2* and *ggpubr*. The Venn diagram was visualized with the help of g*gVennDiagram* (Gao and Dusa, 2024).

Cytosine methylation levels were calculated as a proportion of methylated over total reads. PCA was performed on the whole genome bisulfite sequencing data using the *factoextra* package (Kassambra and Mundt, 2020). For this, all methylation levels were averaged out on 100 bp bins by cytosine context. Raw methylation levels were plotted out for the regions encompassing 1000 bp upstream and downstream of the *COL7* (AT1G73870), *COL8* (AT1G49130) and *NF-YA2* (AT3G05690) genes, according to the TAIR10 assembly, using *ggplot2* and *ggpubr*. Statistical significance for differential methylation was estimated with a mixed linear model (*lme4*) evaluating treatment, genotype and their interactions, as well as the block structure as a random effect. For the search of differentially methylated loci (DML), this was performed for every single cytosine. A rolling window approach was used for the location of differentially methylated regions (DMR), where the mean methylation for 100 bp was estimated and used for the statistical analysis on per cytosine context basis.

## Results

### Impact of water deficit on flowering time

To evaluate the interaction between cytosine methylation and water deficit on flowering time in *Arabidopsis thaliana*, a total of 160 plants were grown in a randomized complete block design (Figure 1). The control plants were watered twice a week, while drought was applied as a lack of water for two weeks, starting at 12 days after seeding (DAS). 96 plants were sampled at 24 DAS for examination. At this stage, water deficiency had reduced the rosette biomass of Col-0 by approx. 60%, while the reduction in *ddc* was closer to approx. 50% (Figure 2A). While *ddc* naturally produces less biomass than Col-0, water deficit reduced the difference between the two genotypes by half. Very similar pattern was observed for rosette diameter (Figure 2B), with the significant difference of *ddc* being closer to Col-0 in diameter than in weight. This led us to hypothesize that *ddc* loses less water under drought than Col-0, leading to less weight loss and less water stress. To test this hypothesis, relative water content (RWC) was measured from leaf number 5 at 24 DAS from 85 plants. Surprisingly, we found that there is no significant difference between Col-0 and *ddc* in both control and low watering treatments (Figure 2C). Leaf number 5 is an already relatively old leaf at 24 DAS, and in some cases exhibits symptoms of senescence. This led to a relatively high biological variation as visible on the graph (Figure 2C). Nevertheless, the visual assessment of the plants also indicated no wilting of the leaves, despite the significant reduction in plant growth. Significant reduction in plant growth was also combined with delayed bolting in Col-0, with an average of 1 day (Figure 2D). No impact on flowering time was observed in *ddc*, which was already exhibiting a delay between 1 and 2 days compared to Col-0 under control conditions. This indicates that the pathways responsible for delaying bolting in Col-0 may already be active in *ddc*. It is interesting to note, that in both *ddc* and Col-0, the water deficit led to a more uniform time of bolting for all plants, essentially reducing the number of both early and late flowering individuals.

**Figure 2.**
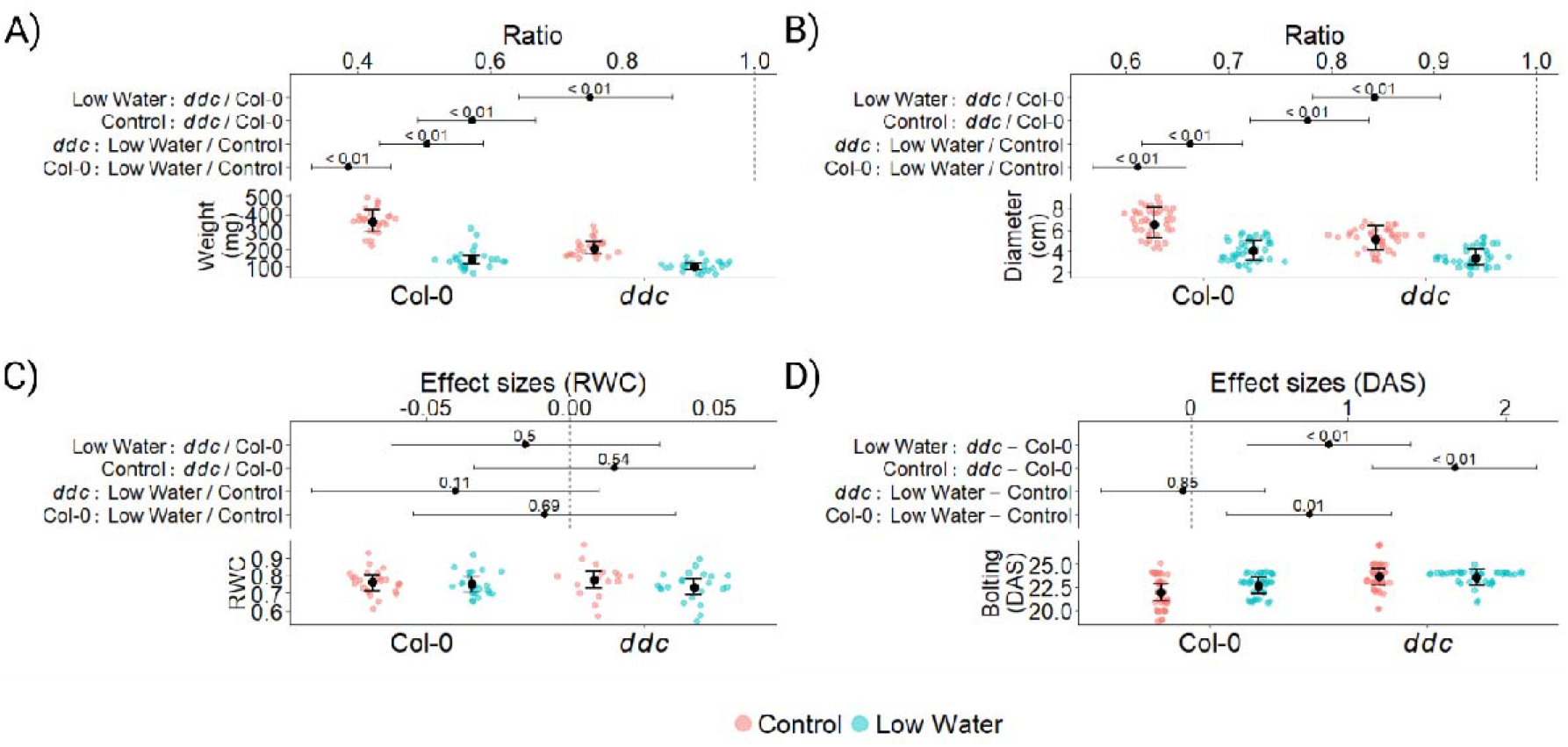
Plant growth and phenotype of *Arabidopsis thaliana* wild type Col-0 and *ddc* triple knockout mutant under drought stress. The data was analyzed using mixed linear models. Red dots represent the Control treatment, blue dots represent the Low Watering treatment. Black dots are the estimated marginal means with whiskers representing the 95% CI. The numbers on the graph represent *p values* of the differences produced by the linear models. Logarithmic transformation was applied to the weight and rosette diameter for better model fit. Therefore the resulting effect sizes are in the form of ratios. A) Plant weight at 24 DAS, measured in mg; n = 96. B) Rosette diameter at 24 DAS, measured in cm; n = 160. C) Relative water content (RWC) at 24 DAS; n = 85. D) Time of Bolting measured in days after seeding (DAS); n = 160.

### Changes in gene expression under drought stress

RNAseq was performed in order to test the hypothesis that a commonly up-regulated pathway between *ddc* under control conditions and Col-0 under Low Water treatment is responsible for the delay in bolting. The principle component analysis shows independent separation between the four treatment groups (Figure 3A), with PC1 separating Control from Low Water treatments and PC2 separating Col-0 from *ddc* genotypes. We found 5239 differentially expressed genes in Col-0 under water deficit (Figure 3B; Supplementary Table 3), compared to 3307 in *ddc* (Figuure 3D; Supplementary Table 5). This indicates that *ddc* might be experiencing less severe stress, as shown by less growth reduction (Figure 2A and B). On the other hand, some pathways which are differentially regulated in *ddc* due to the mutation background may be influenced by drought stress in Col-0. In fact there are 1356 differentially expressed genes in *ddc* compared to Col-0 under control conditions, with the majority being up regulated genes (Figure 3C; Supplementary Table 4).

**Figure 3.**
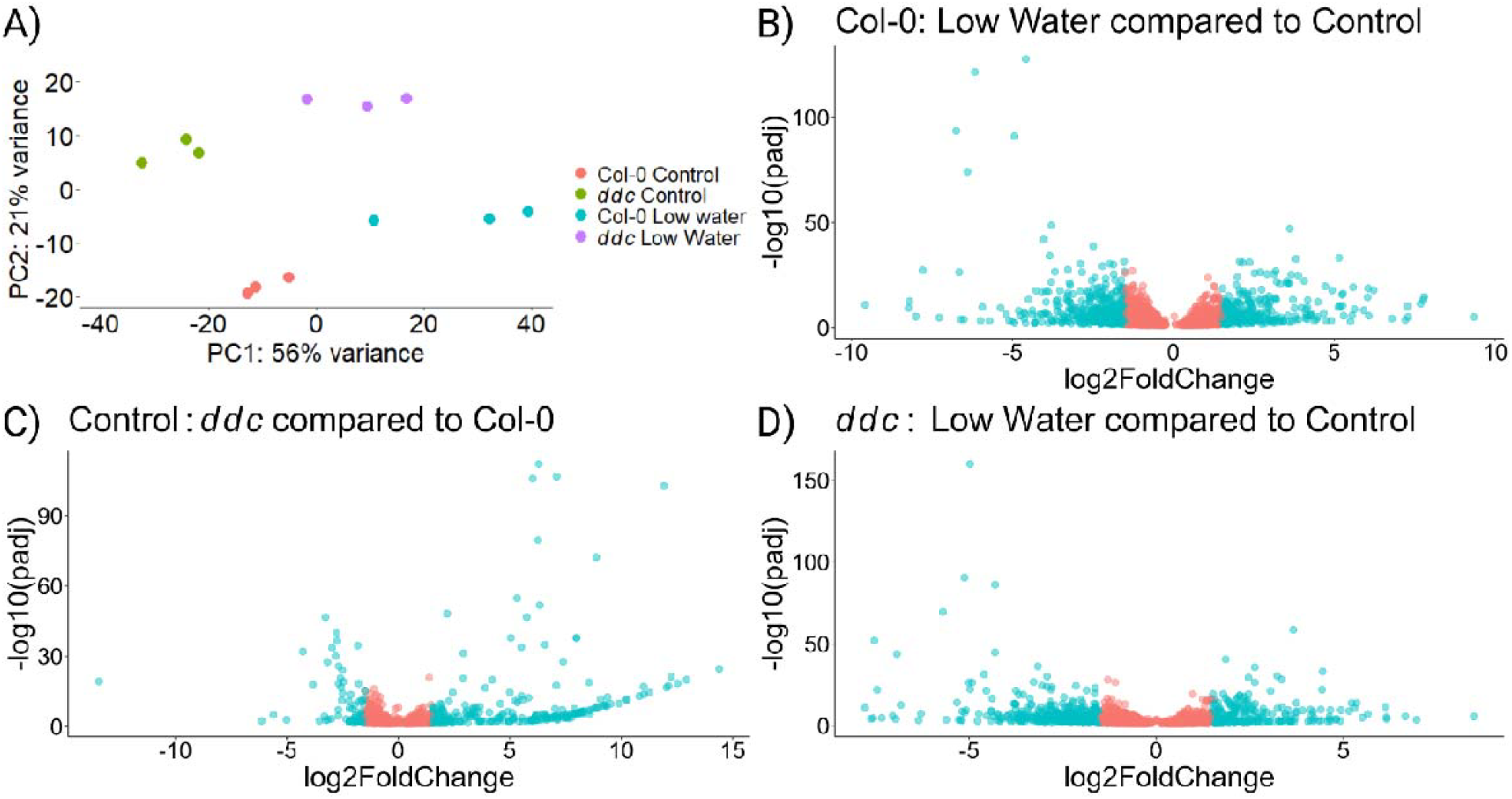
Gene expression of *Arabidopsis thaliana* wild type Col-0 and *ddc* triple knockout mutant under drought stress. 48 plant samples were pooled in 4 groups with 3 biological replicates each. A) PCA analysis performed on the log transformed gene expression data. B) Volcano plot of differentially expressed genes under Low Water compared to Control conditions in Col-0 wild type; n = 5239. B) Volcano plot of differentially expressed genes in *ddc* compared to Col-0 under control conditions; n = 1356. C) Volcano plot of differentially expressed genes under Low Water compared to Control conditions in *ddc* triple knockout mutant; n = 3307. The genes represented on the volcano plots were selected with *padj* < 0.05. Red colored dots represent genes with log2FoldChange < 1.5, blue colored dots represent genes with log2FoldChange >= 1.5.

Next we wanted to find genes whose patterns of expression matches the patterns of bolting observed on Figure 2D. First, differentially expressed genes in Col-0 under Low Water compared to Control treatments were intersected with *ddc* compared to Col-0 under Control (Figure 4A). From these, only genes that are commonly up- or down-regulated were chosen, producing a set of 253 genes. Finally, all genes that are differentially expressed in *ddc* under Low Water treatment compared to control were removed, leaving 188 genes. No significant enrichment was found regarding molecular function, or biological process when probing The Gene Ontology Resource (Ashburner et al., 2000; Aleksander et al., 2023). Looking at cellular components: 67 were chloroplast related, with 71 related to plastids overall. Only three of the 188 genes were previously recognized regulators of flowering. These are the B-box type zinc finger proteins *CONSTANS-LIKE 7 (COL7/BBX16), CONSTANS-LIKE 8 (COL8/BBX17)* and the *NF-YA2* transcription factor (Figure 4B). Another gene was found, which showed suppressed gene expression, but was also significantly down-regulated in *ddc* under Low Water treatment. This gene encodes the DELLA protein *RGA-LIKE1 (RGL1)* (Suplementary Table 6).

**Figure 4.**
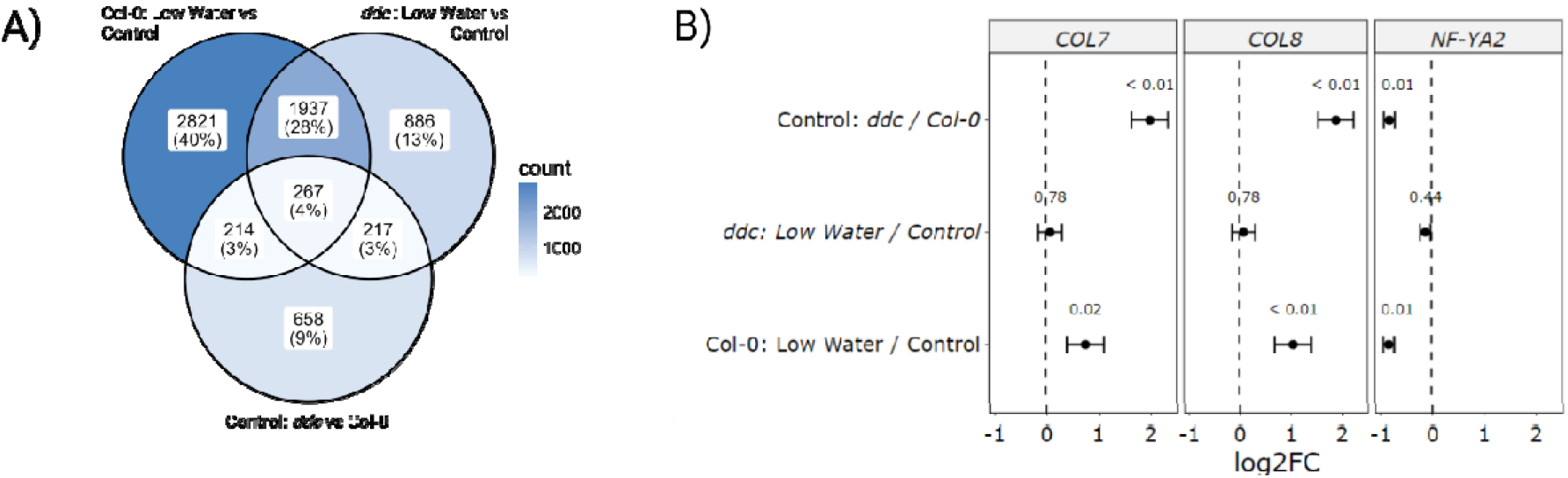
Gene expression analysis of Col-0 wild type and *ddc* triple knockout mutant under drought stress. A) Venn diagram of all differentially expressed genes in three pairwise comparisons: Col-0: Low Water vs Control, *ddc*: Low Water vs Control, Control: *ddc* vs Col-0. B) Differential expression of *COL7, COL8* and *NF-YA2* in three pairwise comparisons. Black dots represent the mean log2 Fold Change (log2FC), whiskers represent the standard errors of the log2FC, and the small numbers above the means represent the adjusted *p* values.

To find alternative explanation for the changes in flowering time, we scanned the three pairwise comparisons of interest for differentially expressed genes related to flowering (Supplementary Table 6). In *ddc* compared to Col-0 under Control, *CCA1* was down-regulated, which also corresponded to induced expression of *TOC1*. Surprisinlgy, C*CA1* was up-regulated in both Col-0 and *ddc* under Low Water, indicating that down-regulation of *CCA1* is a methylation specific response, which is reversed by water deficit. This however, did not appear as a respective down-regulation of *TOC1*. Additionally, Low Watering enhanced the expression of *AP1, SNZ* and Erecta and suppressed the expression of *AP2* in both Col-0 and *ddc*. Overall, 19 genes were differentially expressed in Col-0 under Low Water, compared to 9 genes in *ddc*. Only 7 genes were differentially expressed in *ddc* compared to Col-0 under Control. These results underline the significant role *DDC* induced cytosine methylation plays in regulating flowering time under water deficit.

### Changes in cytosine methylation under water deficit

To establish the role of cytosine methylation in the regulation of genes involved in bolting, whole genome bisulfite sequencing was utilized. The principle component analysis (PCA) shows little effect of the Low Watering treatment on the cytosine methylation landscape (Figure 5A). The two large differences are between Col-0 and *ddc*, separated by PC1. PC2 shows separation of the data points mainly based on the treatment differences and the internal variation between the samples. While Col-0 samples become somewhat separated, *ddc* shows significant overlap between the two groups (Figure 5A). The statistical evaluation shows that there is an overall increase of methylation in all three cytosine contexts (Figure 5B, C and D). CG methylation shows an increase, which is non-significant in Col-0 and close to significance in *ddc* (Figure 5B). Both CHG and CHH methylation show significant increases in methylation levels following Low Watering (Figure 5C and D). These results indicate that *de novo* methylation as a response to the applied stress in the CG and CHH contexts are mostly *DDC* independent. On the other hand, CHG *de novo* methylation is highly reduced, but not lost in the *ddc* mutant. The mean CHG methylation for *ddc* is approx. 1.2% under Low Water and 1.1% under control conditions. The mean CHG methylation for Col-0 is approx. 8.5% under Low Water and 8% under control conditions. This indicates that approx. 20% of the CHG *de novo* methylation in this experiment, is *DDC* independent.

**Figure 5.**
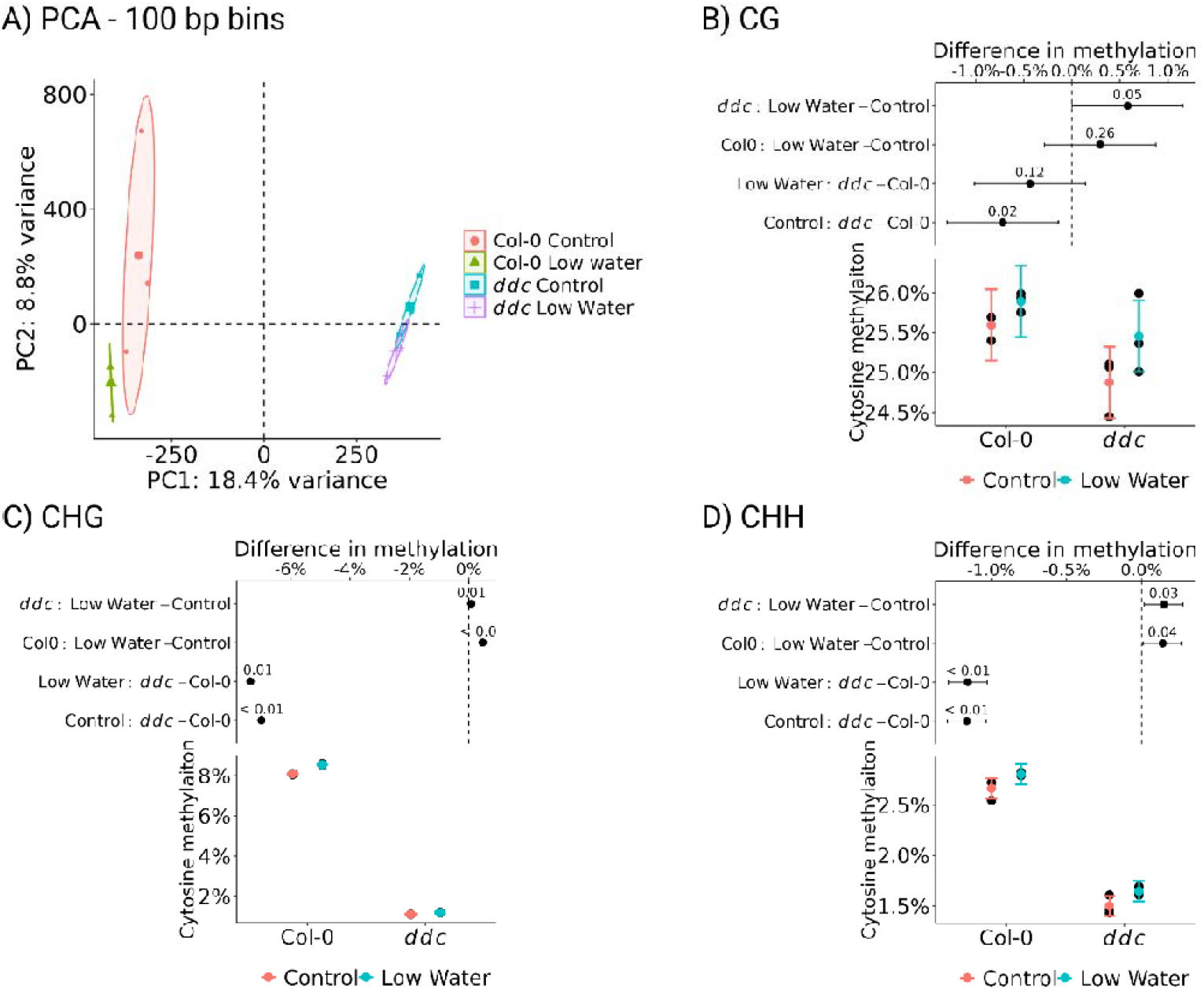
Whole genome bisulfite sequencing data of Col-0 wild type and *ddc* triple knockout mutant under drought stress. 48 plant samples were pooled in 4 groups with 3 biological replicates each. A) PCA on the methylation levels of all cytosine contexts averaged out using 100 bp bins. Mean cytosine methylation of the B) CG, C) CHG and D) CHH contexts are presented as estimated marginal means with 95% CI (bottom); and mean difference levels with 95% CI and *p* value of the mean difference (top).

Next, the cytosine methylation levels were plotted for *COL7* and *COL8*, within 1000 base pairs from the gene loci, including the 3’UTR and the 5’UTR, according to the TAIR10 assembly (Figure 6). COL7 shows existence of cytosine methylation in all three contexts, while *COL8* is methylated mostly at CHG and CHH. No differentially methylated loci, or regions were found related to the two genes that could explain the observed changes in expression patterns (Supplementary Figures 5 and 6). It is also interesting to note that *ddc* triple knockout leads to little, if any change in the methylation landscape, with the exception of a significant increase in CG methylation towards the 3’UTR of *COL8*. These results indicate that methylation in these two genes is established and maintained in a manner independent of the *DDC* methyltransferases. Furthermore, the increase in gene expression resulting from the *ddc* mutations is likely not a direct cause of changes in methylation in cis, but rather is a result of siRNA/miRNA production, or changes in certain signal transduction pathways that impact the two genes in trans. Thus, this experiment shows that cytosine methylation plays a role in fine tuning flowering time under water deficiency, potentially through regulating the expression of the *COL7* and *COL8* zinc finger proteins in trans. Further research is required to establish the exact mechanism by which *COL7* and *COL8* are regulated and the significance of their impact on flowering time under different stressful environments.

**Figure 6.**
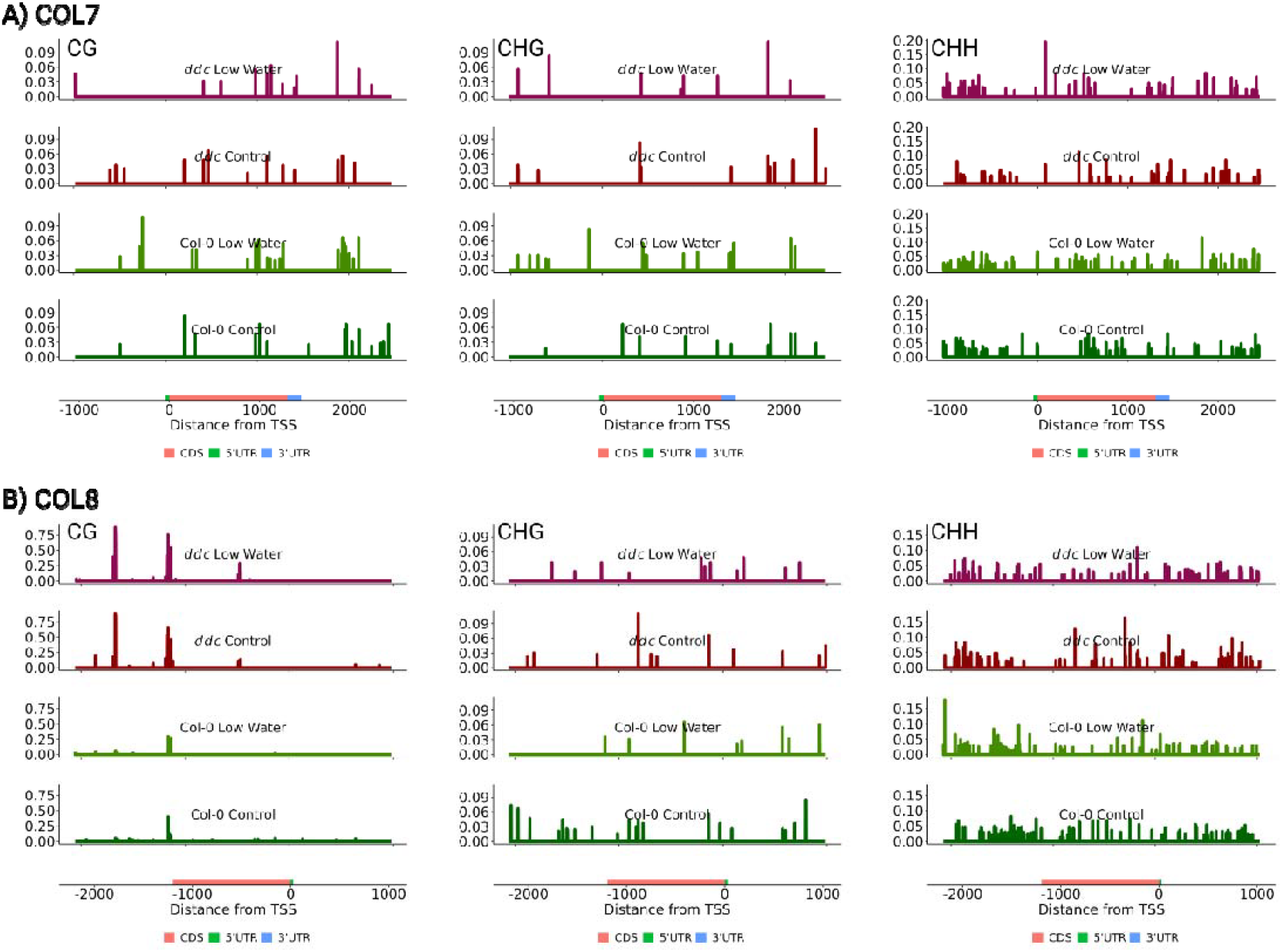
Mean cytosine methylation levels from three biological replicated of CG (left), CHG (middle) and CHH (right) contexts for A) *COL7* and B) *COL8*. On the x axis, transcription start site is labeled as 0 with position numbers increasing according to the positive DNA strand. Y axis measures cytosine methylation proportion for each individual cytosine. The four experimental conditions are presented from top to bottom as follows: *ddc* Low Water, *ddc* Control, Col-0 Low Water and Col-0 Control.

While not much is known about the regulation of COL7 and *COL8* and their role in stress induced changes of flowering time, *NF-YA2* is a relatively well characterized gene. Under drought stress, *NF-YA2* is downregulated via *miR169d* (Gupta et al., 2024). First, we mapped the methylation profile of the *NF-YA2* gene together with 1000 base pairs up and down stream of the gene (Figure 7). No obvious changes in the methylation profile were found, that could explain the expression under water deficit, or in the *ddc* mutant. The gene appears to have CHG and CHH methylation levels comparable to those of *COL7* and *COL8* (Figure 6), again without much impact of the *ddc* mutations. Surprisingly, we found two CG methylation peaks towards the 3’UTR end of the gene and downstream of the *miR169d* target region. This region gained additional CG methylation in the *ddc* mutant (Supplementary Figure 3). Within the target area of *miR169d* however, significant gain of CG methylation was observed, but only for the *ddc* mutant and irrespective of the treatment (Figure 7; Supplementary Figures 3 and 4). A slight reduction in CHG was also observed under water deficit in Col-0 and complete loss in *ddc*, but this was not deemed significant by the statistical model. Additionally, an increase in CHH methylation was observed downstream of the *miR169d* target region, which appeared in Col-0 under low water and *ddc* under control, but was lost in *ddc* under Low Water. Whether these effects are stochastic in nature remains to be elucidated. The gene expression patterns and the phenotype in the present experiment suggest that these changes in cytosine methylation are not a cause but rather an effect of the interaction between *miR169d* and the *ddc* triple knockout background.

**Figure 7.**
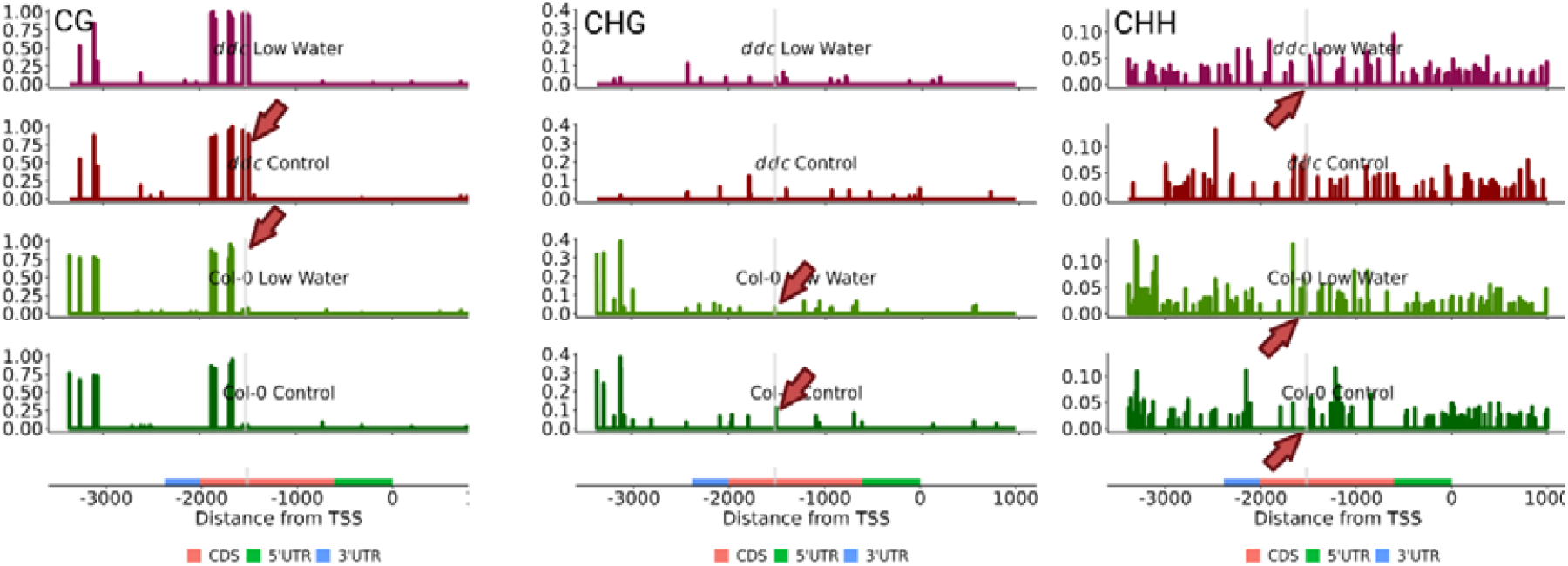
Mean cytosine methylation levels from three biological replicates of CG (left), CHG (middle) and CHH (right) contexts for *NF-YA2*. The four experimental conditions are presented from top to bottom as follows: *ddc* Low Water, *ddc* Control, Col-0 Low Water and Col-0 Control. Red arrows point at the location of *mir169d* interaction, where changes in the cytosine methylation profile are observed. Light grey lines indicate the target site of *mir169d*.

## Discussion

There are two general strategies *Arabidopsis thaliana* plants utilize in order to cope with drought stress – drought escape and draught tolerance. The wild type Col-0, which is often used in laboratory experiments, is a drought escape accession with a nonfunctional *FRI* allele resulting in earlier flowering (Lovell et al., 2013). We observed that, in its growth pattern under water deficiency, Col-0 does not lose turgor for a long period of time (Figure 2C; Bouchabke et al., 2008). Rather, it slows down growth and delays further development (Figure 2A and B). Transition to flowering still occurs but with a delay (Figure 2D). When the water deficit becomes critical, the plants lose turgor rapidly and die.

We had previously observed that the *ddc* (*drm1/drm1/cmt3*) triple knockout mutant shows a slight delay in its transition to flowering under long day conditions (Vatov et al., 2022). In the present study we tested if the delay from the *ddc* mutations is additive to that from water deficit and found that it is not (Figure 2D). The first obvious gene candidate for this effect is the *FLC*. It has been long known that *FLC* is under trans acting control from DNA methylation, but the exact mechanism remains to be elucidated (Finnegan et al., 2005; Shi et al., 2023). Nevertheless, hypomethylation is generally related to *FLC* downregulation (Finnegan et al., 2005). On the other hand, drought stress was shown to induce *FLC* expression in an ABA dependent manner (Wang et al., 2013). Indeed, we found that *FLC* is significantly up regulated in the Col-0 wild type under water deficit, but not in *ddc* Control, or *ddc* Low Water (Supplementary Table 6). It is important to note that there is no significant difference between *FLC* expression in Col-0 Low Water and *ddc* Low Water, indicating small increases in expression in *ddc* compared to Col-0 under Control conditions, and *ddc* Low Water compared to Control, which are not considered significant by the statistical model.

In *ddc* compared to Col-0 under control conditions, we found the slight, but significant overexpression of *SOC1* (Supplementary Table 6). This gene is long known as a central regulatory hub for induction of flowering in *Arabidopsis* together with *FT* (Samach et al., 2000). Another surprising find in this comparison is the up regulation of *TOC1* and the corresponding down regulation of *CCA1* (Supplementary Table 6). *TOC1* is generally recognized as a stabilizer of *CO* and promoter of flowering, while *CCA1* itself is a negative regulator of *FT* and *SOC1* (Nakamichi et al., 2007; Lu et al., 2012; Park et al., 2016; Yang et al., 2024). Additionally *CCA1* appears to be up regulated in both *ddc* and Col-0 under water deficit, indicating that cytosine methylation dependent and stress related gene regulation act via two distinct pathways. *CCA1* up regulation could in fact correspond to delayed flowering time under Low Water, however the lack of additional effect in *ddc* indicates another pathway which overrules the effects of *CCA1*.

Another part of the regulation of flowering time recognized in this study is the *NF-YA2* gene, part of the *NF-Y* conserved transcriptional regulator complex. Previous reports suggest that *NF-YA2* promotes *FLC* expression (Xu et al., 2014; Chen et al., 2023) and therefore suppresses the transition to reproductive growth. In our experiment *NF-YA2* expression is down-regulated in Col-0 under water deficit and in *ddc* in control conditions and water deficit. However, this does not match the expression patterns of *FLC*. On the other hand, evidence exists that *NF-YA2* promotes expression of *FT* via direct interaction with the *FT* promoter (Siriwardana et al., 2016). *FT* expression in the present study is not differentially regulated in any of the conditions. This is most likely due to harvesting of the plant material several hours after the beginning of their day, resulting in low base reads from the *FT* gene (Supplementary Table 6).

Recent studies brought insight into the control of *NF-YA2* under drought and heat stress (Gupta et al., 2024). The researchers demonstrated that overexpression of *miR169d* under drought stress resulted in down regulation of the *NF-YA2* gene. We found that down regulation of the *NF-YA2* gene is additionally correlated with some interesting changes in the cytosine methylation landscape, within and downstream of the location targeted by *miR169* (Figure 7; Supplementary Figures 3 and 4). The appearance of CG methylation peaks in methylation deficient mutants, including *ddc*, was described as early as 2005 (Tran et al., 2005). Here we observe that such a peak can appear as a result of the action of miRNA silencing in the *ddc* background. The mechanisms behind this remain to be elucidate.

Finally, two b-box type zinc finger proteins were found to be overexpressed in Col-0 under water deficit and in *ddc* under control. These are the *BBX16* and *BBX17* otherwise known as *CONSTANS-LIKE 7 (COL7)* and *CONSTANS-LIKE 8 (COL8)* respectively. Recently, *BBX16/COL7* was recognized as a regulator of flowering time via direct interaction with the CO protein in the nucleus (Susila et al., 2023). The BBX-CO interaction significantly reduced the ability of CO to induce *FT* expression and was therefore recognized as a negative regulator of flowering time. Additionally, overexpression of *BBX16/COL7* resulted in delayed flowering in *Arabidopsis*. One year earlier it was shown that *BBX17/COL8* has a similar function of interacting with the CO protein and inhibiting its activity (Xu et al., 2022). To our knowledge, there is no known data showing the involvement of these two genes in flowering regulation under water deficit.

The cytosine methylation data does not show any obvious alterations that could cause the differential expression patterns observed in this study. The significant increase in CG methylation towards the 3’ end of *COL8* however, resembles the increase observed in *NF-YA2*. Additionally, there is an inconsistency between TAIR10 (used in this study) and the the Araport11 assembles. Araport11 indicates that the *COL8* transcript expands approx. 400 bp longer as a 3’UTR towards the second *ddc* dependent CG methylation peak (Figure 6). This observation poses the question of the potential regulation of *COL8* via an unknown miRNA pathway. The proof of the CG hypermethylation peaks resulting from the activity of miRNAs in the *ddc* background however, is outside of the scope of this manuscript.

## Conclusion

The results in this experiment support a model of delayed flowering in Col-0 under water deficit by inhibition of the *FT* inducers *CO* and *NF-YA2*. Under long day conditions, the BBX16/COL7 and BBX17/*COL8* proteins interact with CO to inhibit its function as *FT* promoter (Figure 8). On the other hand *miR169d* overexpression down regulates *NF-YA2*, which is in turn an independent promoter of *FT*. Col-0 contains a non-functional *FRI* allele and therefore is unable to induce significant inhibition of the *FT* gene via *FLC*. This results in suppression of the transition to reproductive growth, which however can be easily overcome. The end effect is the slight delay in flowering time observed in this experiment. No additional delay in flowering time in *ddc*, together with matching patterns of expression of the three genes, indicate the significant role of cytosine methylation in this process. How exactly does cytosine methylation impact the expression of *BBX16/COL7, BBX17/COL8* and *NF-YA2* in *trans*, remains to be elucidated.

**Figure 8.**
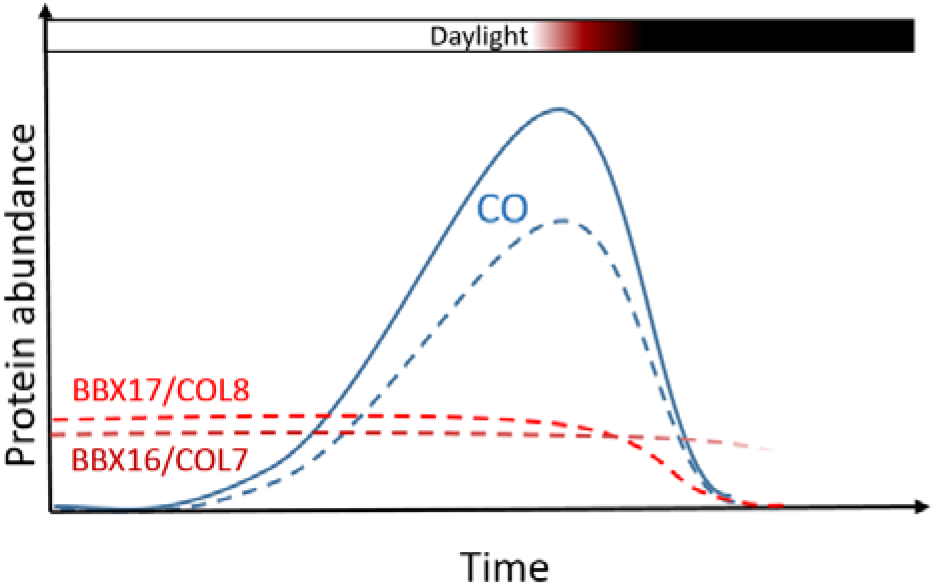
Under normal conditions, the expression of *CO* peaks during the late hours of the day, leading to the accumulation of the CO protein. During the night the expression of *CO* is inhibited, while its protein product gets actively degraded. Long days lead to induction of flowering due to the accumulation of *CO* before sunset (solid blue line), which in turn induces the expression of the florigen *FLOWERING T (FT)*. Drought stress induces the expression of the *BBX16/COL7* (dashed dark red line) and *BBX17/COL8* (dashed red line) genes, whose products interact with CO and inhibit its function as *FT* inducer (dashed blue line), leading to delays in bolting. It is known that *BBX17/COL8* gets actively degraded during the night in a *COP1*-dependent manner. The daily patterns of expression and product accumulation of *BBX16/COL7* and *BBX17/COL8* remain to be elucidated.

## Acknowledgements

This work was supported by the Bulgarian National Science Fund, Petar Beron program (Project EpiFlowScen, grant No. КП-06-ДБ/2), the European Union’s Horizon 2020 research and innovation program, project PlantaSYST (SGA-CSA No. 739582 under FPA No. 664620), and the BG05M2OP001-1.003-0001-C01 project, financed by the European Regional Development Fund through the Bulgarian “Science and Education for Smart Growth” Operational Program, and the Program for Research, Innovation and Digitalisation for Smart Transformation (PRIDST) Operational Programme.

